# *In vivo* intraoral waterflow quantification reveals hidden mechanisms of suction feeding in fish

**DOI:** 10.1101/2021.09.17.460802

**Authors:** Pauline Provini, Alexandre Brunet, Andréa Filippo, Sam Van Wassenbergh

## Abstract

Virtually all fish rely on flows of water to transport food to the back of their pharynx. While external flows that draw food into the mouth are well described, how intra-oral water flows manage to deposit food at the esophagus entrance remains unknown. In theory, the posteriorly moving water must, at some point, curve laterally and/or ventrally to exit through the gill slits. Such flows would eventually carry food away from the esophagus instead of towards it. This apparent paradox calls for a filtration mechanism to deviate food from the suction-feeding streamlines. To study this gap in our fundamental understanding of how fish feed, we developed and applied a new technique to quantify three-dimensional patterns of intra-oral water flows *in vivo.* We combined stereoscopic high-speed x-ray videos to quantify skeletal motion (XROMM) with 3D x-ray particle tracking (XPT) of approximately neutrally buoyant spheres of 1.4 mm in diameter. We showed, for carp (*Cyprinus carpio*) and tilapia (*Oreochromis niloticus*), that water tracers displayed higher curvatures than food tracers, indicating an inertia-driven filtration. In addition, tilapia also exhibited a ‘central jet’ flow pattern, which aids in quickly carrying food to the pharyngeal jaw region. When the food was trapped at the branchial basket, it was resuspended and carried more centrally by periodical bidirectional waterflows, synchronized with head-bone motions. By providing a complete picture of the suction-feeding process and revealing fundamental differences in food transport mechanisms among species, this new technique opens a new area of investigation to fully understand how most aquatic vertebrates feed.

## Introduction

Waterflows are essential for transporting food inside the buccal cavity in a large variety of aquatic animals, including virtually all fish ^1–3^. During suction feeding—the process in which a powerful expansion of the buccopharyngeal cavity causes inflow into the mouth ^4^—the three-dimensional (3D) waterflow patterns toward the fish’s mouth have been particularly well studied ^5–8^. Although waterflow patterns external to the mouth depend on the approaching speed of the fish ^9,10^, instantaneous streamline patterns are fairly conserved between species and behaviors ^11^: as the 3D streamlines converge toward the gape (Fig. 1a, blue arrows), food items that are initially somewhere close to the mouth will curve into the buccal cavity by these extraoral waterflows.

**Fig. 1.**
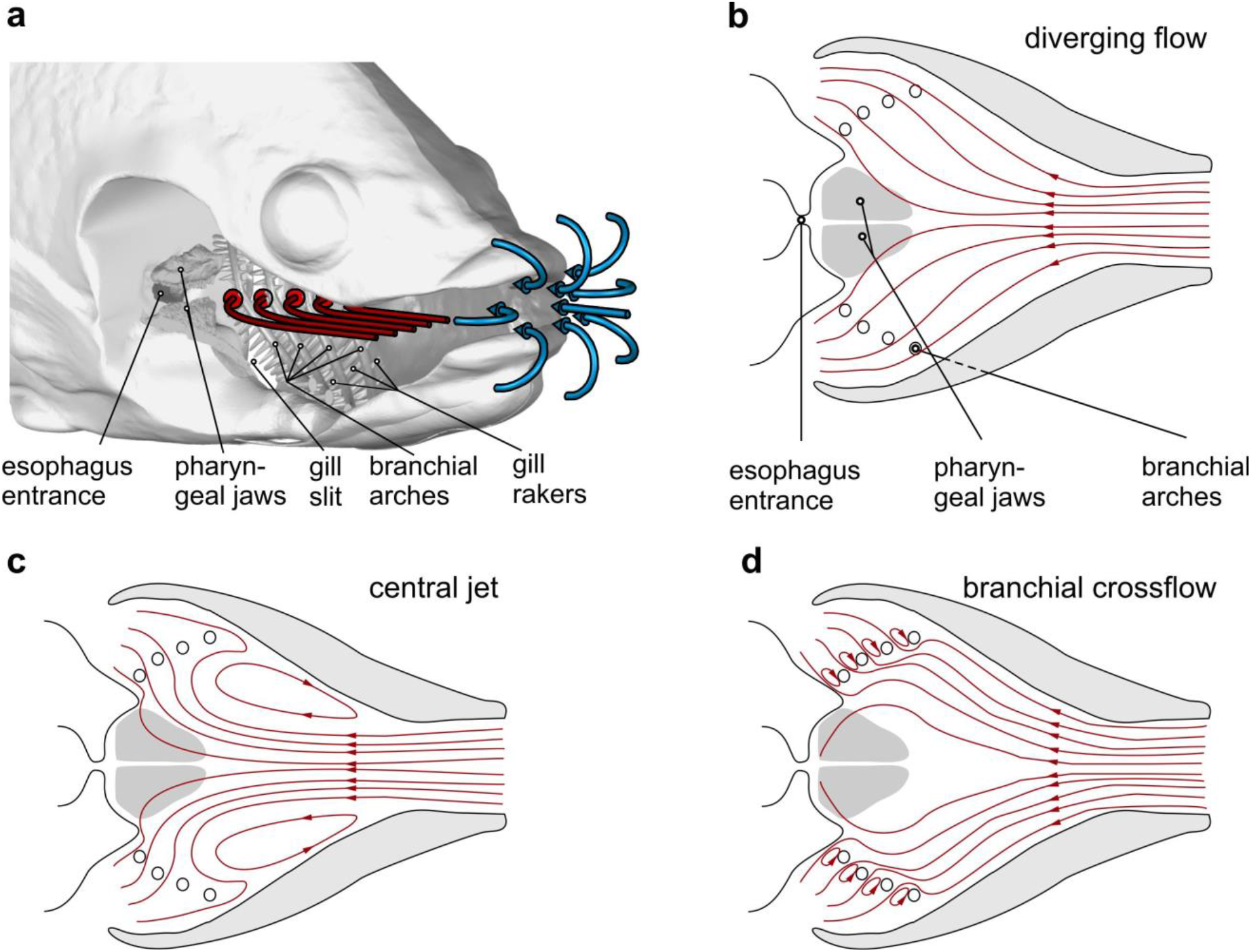
Hypothetical waterflow patterns during suction feeding in fish. a. Schematic illustration of extraoral flows (blue pathline arrows) and hypothetical intraoral flows (red pathline arrows) with important anatomical structures involved in food interception (the right-side buccopharyngeal walls and branchial basket are removed) in a bony fish. **b-d**. Three hypotheses for intra-oral flow patterns, viewed in a mid-frontal section plane. Note that from **b to d**, the tendency to move food away from the desired food target, the esophagus, decreases.

In contrast to this relatively straightforward and well-resolved mechanism of food capture toward the mouth, food transport mechanisms inside the buccopharyngeal cavity are not as simple ^12–15^ and are largely unknown ^16^. After passing through the mouth opening (Fig. 1a, blue arrows), water streamlines are generally assumed to diverge along with the widening buccopharynx (Fig. 1a,b, red arrows) ^17,18^. This implies that, inside the buccal cavity, waterflow tends to be directed toward the gill slits located at the posteroventral and posterolateral head margins (Fig. 1b). Paradoxically, as these flows take a short route to the gill slits, they seem far from optimal to carry food to the entrance of the esophagus along the midsagittal plane of the buccal cavity.

To date, there is no evidence that the esophagus expands in concert with buccopharyngeal suction generation to directly draw in a significant amount of water together with the food. Therefore, other structures need to be involved in separating food from the waterflows. Ideally, in bony fish, the food that is sucked into the mouth could be intercepted by the pharyngeal jaws located just anterior to the esophageal sphincter (Fig. 1). These mobile pharyngeal jaws, while maintaining physical contact with the food, can move it to the esophagus ^19,20^. However, given the current hypothesis of diverging streamlines in the expanding buccopharyngeal cavity (Fig. 1a,b), food is more likely to end up being sieved by the other branchial arches and their gill rakers, after which a second food transport cycle is needed which encounters the same problem regarding how the food is able to reach the esophagus. Consequently, intraoral hydrodynamics are critical in determining the initial food deposit site after capture and, in turn, the associated anatomical adaptations and behaviors for food transport and handling inside the head.

Alternatively, two other intraoral flow patterns are theoretically possible, which may facilitate interception by the pharyngeal jaws (Fig. 1c-d). Computational models of suction-feeding hydrodynamics ^21,22^ show that flow can separate from the expanding buccopharynx to create a centralized jet of suction flow, along with anterior flow at the borders of the buccal cavity forming vortices (Fig. 1c). This pattern, to which we will refer as a ‘central jet’, could carry food closer to the pharyngeal jaws (Fig. 1c) compared to the classical hypothesis of ‘diverging flow’ (Fig. 1b). Thirdly, as hypothesised for fishes relying on filtration of smaller food particles from the water ^23,24^, gill arch and raker structures may cause crossflows, in which the main direction of flow medial of the branchial arches is directed towards the esophagus and only the permeate of the branchial filter basket flows out through the gill slits (Fig. 1d).

Despite this critical role during feeding in nearly all fish, the spatiotemporal pattern of intraoral waterflow has still not been quantified *in vivo* due to the difficulty in gaining optical access to the buccopharyngeal cavity. This implies a fundamental gap in our knowledge regarding how more than half of all vertebrates feed. Here, we develop a new technique based on biplanar high-speed X-ray videography to simultaneously quantify the 3D pathlines of both food and intraoral water, as well as skeletal kinematics. We apply this technique to two distantly related omnivorous suction feeders, carp (*Cyprinus carpio*) and tilapia (*Oreochromis niloticus*). We use this data to describe the intraoral waterflow and to determine the intraoral mechanisms of aquatic feeding that have remained hidden thus far.

## Results

### Intraoral kinematics of water tracers and food

#### The periodic bidirectional motion of water tracers

During each feeding sequence, the water tracers moved alternatingly backward and forward, interspersed with phases of stasis (Movie S1, Movie S2, Fig. 2). At the beginning of the sequence, we observed a relatively low posterior motion of the water tracers toward the posterior part of the buccal cavity. As the velocity of the water tracers is calculated in reference to the entrance of the esophagus, this low velocity corresponds to the approaching speed of the fish. It corresponds to 3.9 ± 2.5 cm/s in carps and 3.3 ± 1.3 cm/s in tilapias.

**Fig. 2.**
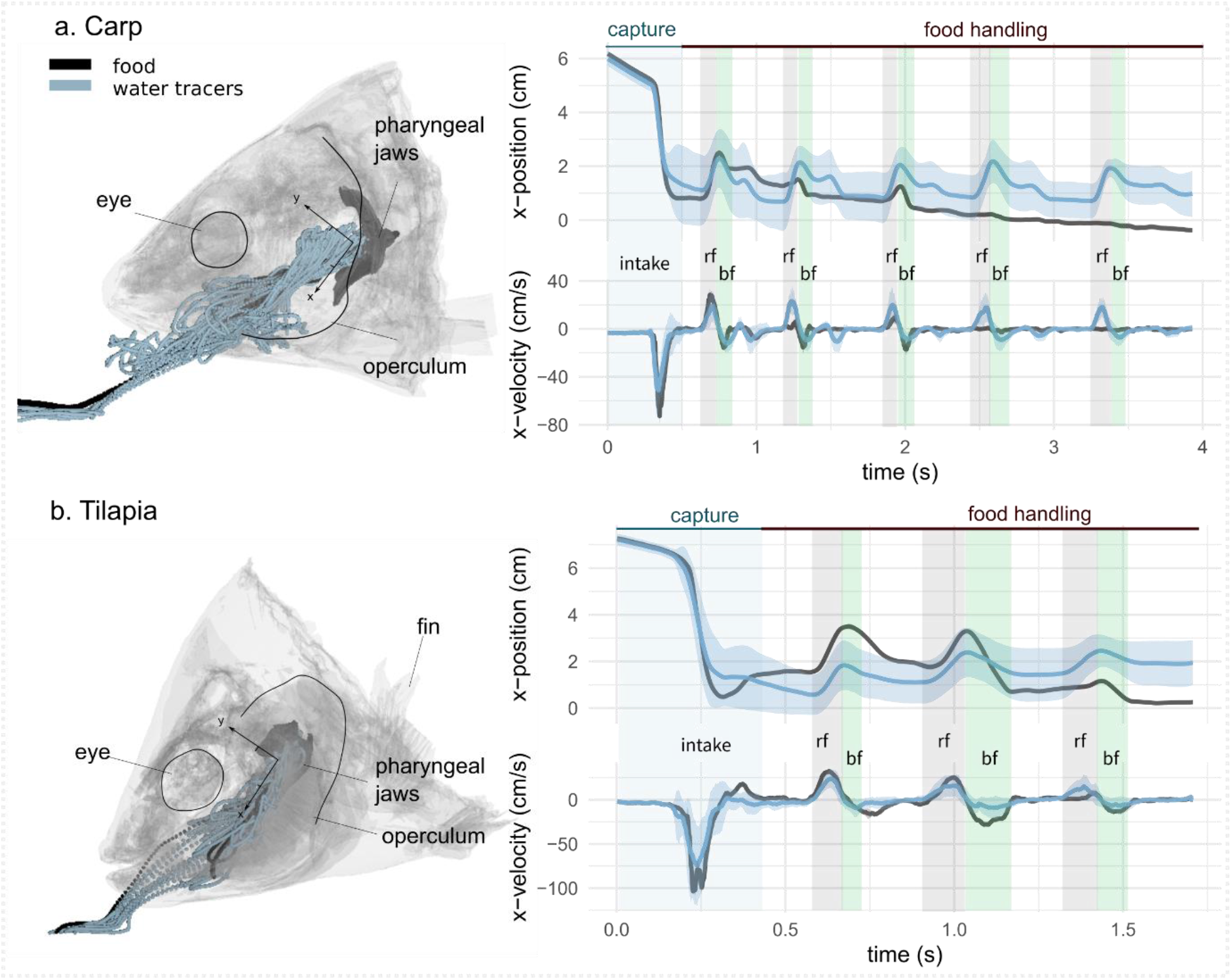
Representative sequences of suction feeding for carp (a) and tilapia (b). Water and food tracer 3D trajectories outside and inside the buccal cavity are represented by a semi-transparent model of a carp (**a**, left) and a tilapia (**b**, left) head. The anteroposterior velocity of the water and food tracers of a carp (**a**, right bottom) and a tilapia (**b**, right bottom) over a representative suction-feeding sequence are shown, with the mean trajectory of the water tracers (blue line) surrounded by the standard deviation (lighter blue), as well as the trajectory of the food tracer (black). On the graphs, we show the division into two main phases: capture and intraoral food handling. During capture, the intake (intake) corresponds to a high negative magnitude for the anteroposterior velocity, followed by stasis, during which the anteroposterior velocity is close to zero. During intraoral food handling, we see an alternance of stasis consisting of a reverse flow with a positive anteroposterior velocity and of a backflow with a negative anteroposterior velocity of lower amplitudes compared to the first strike.

Then, we observed a rapid increase in the posterior velocity of the water tracers, reaching up to 92.1 ± 59 cm/s for the carps and 97.8 ± 23 cm/s for the tilapias. We named this rapid increase “intake” (Fig. 2), corresponding to what is commonly known as the food-capture or food-intake phase.

After the intake, a phase of apparent stasis occurred, with a steady, low-velocity, anteroposterior water tracer trajectory, leading to a nearly complete stop of the food. Then, the water tracers started to move again, which corresponded to the beginning of the intraoral food-handling phase. This phase was characterized by lower velocities compared to the capture phase (the maximum during food capture was 32.9 ± 11 cm/s in carps and 24.7 ± 12 cm/s in tilapias) and by the occurrence of bidirectional motion: the reverse flows (rfs) (previously termed “back-wash”; ^20^), defined by the negative anteroposterior velocities of the water tracers (highlighted in gray in Fig. 2), and backflows (bfs), with a similar pattern as the intake, but slower (highlighted in green in Fig. 2).

The succession of the intake and the alternating rf and bf phases were consistent across trials and could be automatically detected in the studied trials (*N* = 13) (Data file S1). We found that the periodicity of back-and-forth motions (rf and bf) was consistent throughout the sequence. For the carps, the transition between the rf and bf phases happened every 0.58 ± 0.12 s; for the tilapias, it happened every 0.40 ± 0.05 s.

#### Differences between the water tracer and food trajectories

Among the 73 trials recorded, no water tracers passed the esophagus; however, the food was successfully ingested in most cases (5 and 6 failures or spits out of 34 and 36 trials in carps and tilapias, respectively). Among the 7 trials of carps and 6 sequences of tilapia we were able to fully analyze (hydrodynamics and bone kinematics analyses), the total distance traveled by the food tracers during the pooled intake, the rf, and the bf phases was significantly lower than the distance covered by the water tracers (Fig. S2), demonstrating a different path between the food and water tracers.

The 3D radio-opaque particle-tracking method allowed us to export the 3D trajectory of all the water tracers during a suction-feeding sequence. This provided us with a visualization of the water tracer paths for both the lateral and dorsoventral views during the intake (Fig. 3a), the rf phase (Fig. 3b), and the bf phase (Fig. 3c) for a representative trial and for the combination of 3 trials for one individual of carp and 3 trials for one individual of tilapia.

**Fig. 3.**
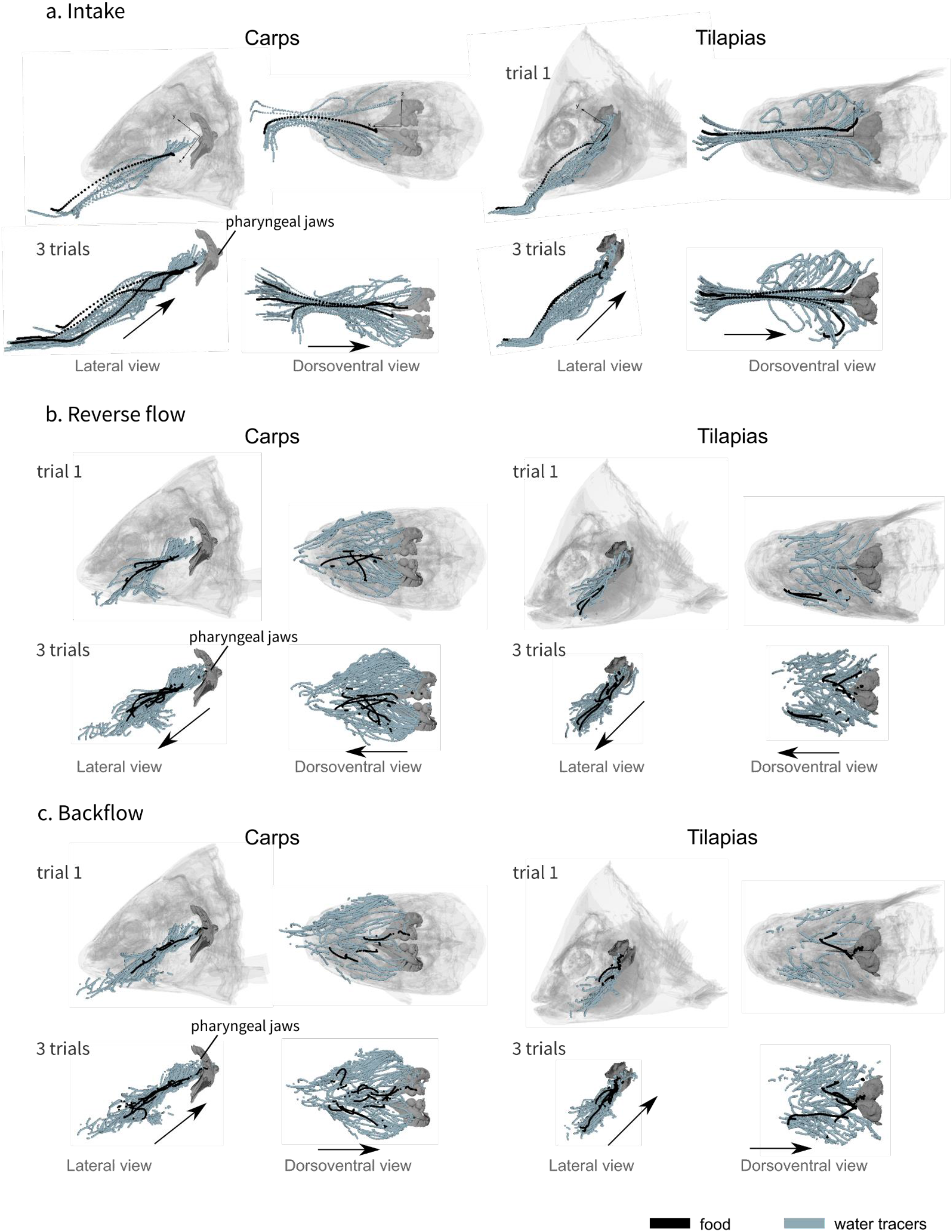
Visualization of the water tracer paths for both the lateral and dorsoventral views during the intake (**a**), the reverse flow (**b**), and the backflow (**c**). A representative trial is presented on the first row of each phase and a combination of 3 trials for one individual of carp and 3 trials for one individual of tilapia on the second row of each phase, showing the consistency of the patterns. The axes are represented on the first row of schematics, the orientation remains the same for all the other schematics. The black arrows represent the global direction of tracer movement.

During the intake, the food item usually followed a trajectory that was more dorsal in comparison to the water tracer trajectory in both species (Fig. 3a). In addition, the food trajectory remained close to the midsagittal plane, whereas the water tracers tended to spread throughout the entire buccal cavity (Fig. 3a). Among the dozens of water tracers captured by the fish during each trial, only 3 to 5 per trial also followed a midsagittal trajectory to reach a position close to the entrance of the esophagus (Fig. 3a). The others spread to the lateral parts of the buccal cavity and stopped at the gill rakers (Fig. 3a). In tilapias, the water tracer deviation started more anteriorly (Fig. 3a), but in both species, the path curvature of the water tracers was significantly higher than that of the food tracers (Fig. 4), demonstrating a more important lateral deviation of the water tracers compared to the food tracers.

**Fig. 4.**
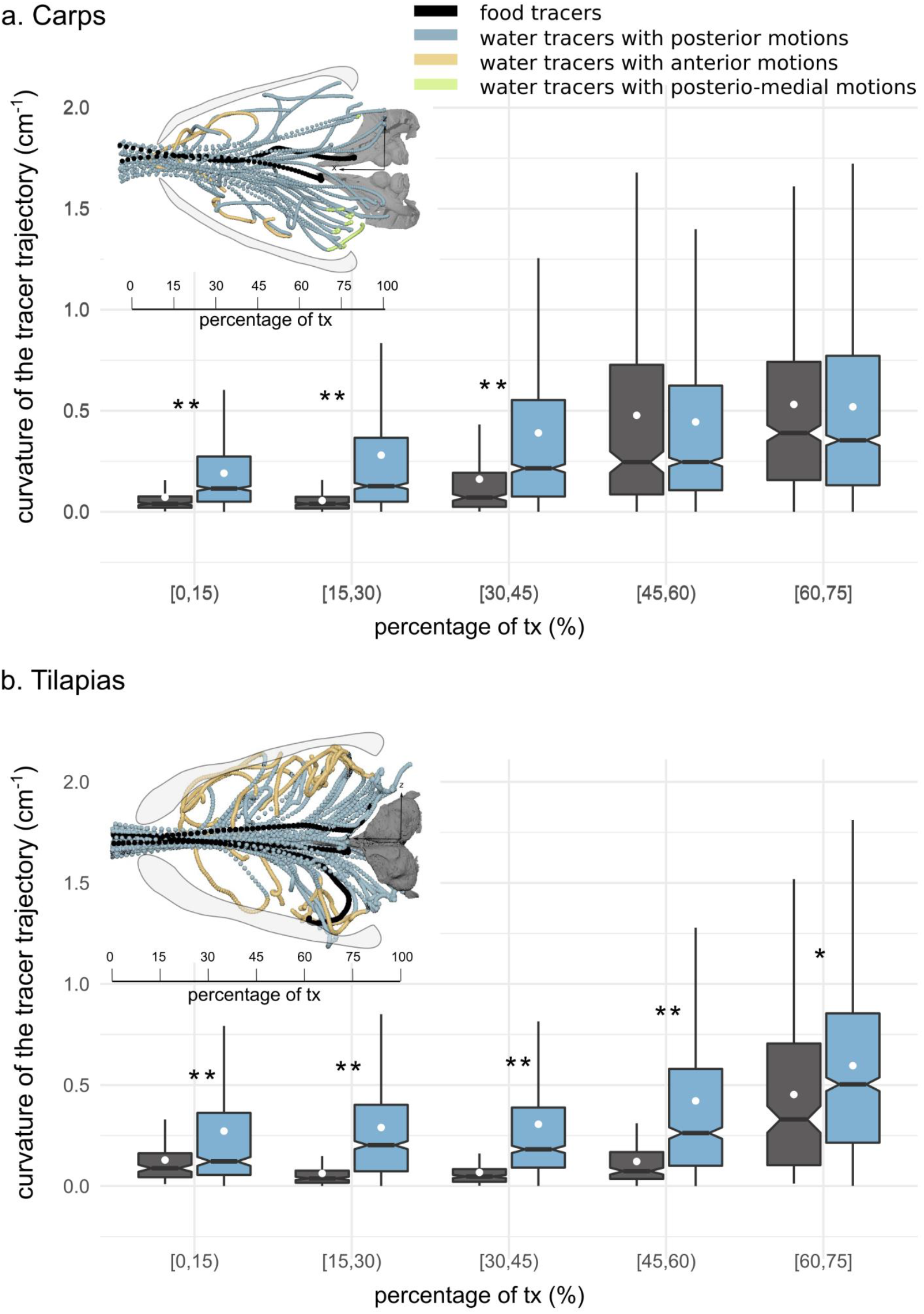
Boxplots of the medio-lateral curvature. calculated based on intervals of the anteroposterior trajectory during the intake in carps (**a**) and in tilapias (**b**). The lower and upper hinges correspond to the first and third quartiles (the 25th and 75th percentiles), and the white dot gives a 95% confidence interval for comparing medians. The asterisks represent the significance of the statistical tests comparing the water and food tracers curvatures (*: P-value ≤ 0.05, **: P-value ≤ 0.01). The schematics in the upper part of each subfigure correspond to a dorsoventral view of 3 trials of one individual of carp (a) and 3 trials of one individual of tilapia (b) showing the water (in blue, yellow and green) and food tracer (in black) trajectories. The blue, yellow and green trajectories highlight the posterior, anterior and posterio-medial motions of the water tracers, respectively.

During the other phases, the water tracers spread throughout the entire buccal cavity, but their trajectory mainly remained anteroposterior in nature (Table S1, Figs. 3b, 3c). After the first part of the intake, there was no apparent correlation between the food and the water tracer trajectories (Figs. 2, 3). This suggests that the food was passively moved by the waterflows, which eventually managed to reposition the food toward the esophagus.

#### Intake flow patterns

Especially for tilapia, many flow tracers move anteriorly after starting to curve in the lateral direction (Fig. 4b). Such anterior flow close to the borders of the buccal cavity is characteristic of the central jet pattern (Fig. 1c). In contrast to tilapia, in carp such flows occur only close to the mouth entrance (Fig. 4a), while flow in the posterior half of the buccal cavity resembles a diverging flow pattern (Fig. 1b). Posterio-medial flows in the branchial arch region, which would be indicative of crossflow (Fig. 1d), could not be discerned. In carp, only one of the many flow tracer paths was directed medially (see green tracers in Fig. 4a). In tilapia, none of the paths showed a clear medial component (Fig. 4b).

### Kinematics of cranial bones

The 3D kinematic analysis of the head bones revealed a stereotypical sequence of motions of the upper and lower jaws (gape), hyoid depression, and opercula abduction.

During the intake (highlighted in blue in Fig. 5), a peak gape was followed by a peak in hyoid depression, followed by a peak in opercula abduction. This sequence of peaks occurred in each trial for both species (Data file S1). The abduction of the opercula is prominent at the time of lateral motion of the water tracers after they enter the buccal cavity.

**Fig. 5.**
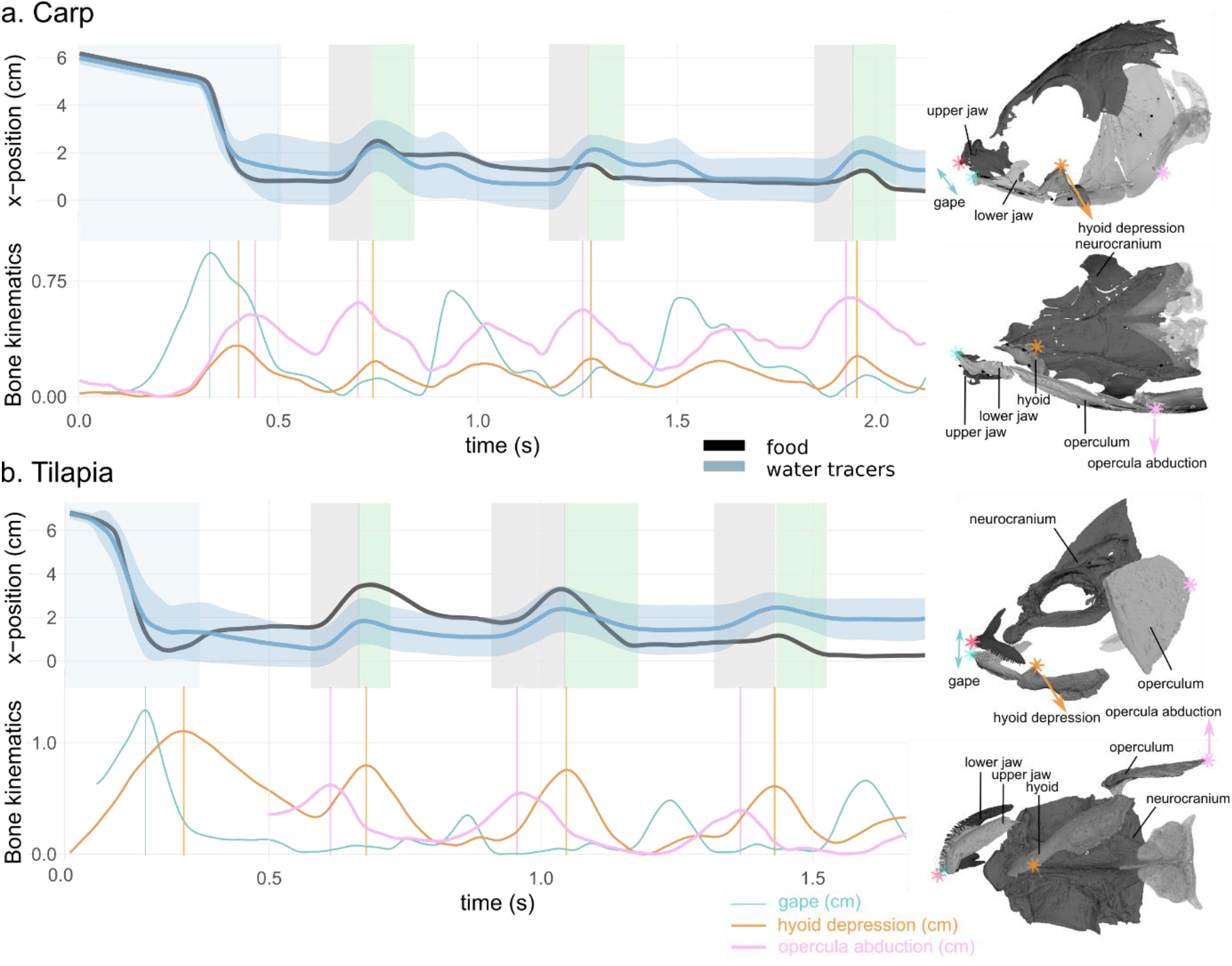
Water tracer anteroposterior trajectory, gape, hyoid depression, and opercula abduction during representative suction-feeding sequences in carp (**a**) and tilapia (**b**). The shaded bars in the x-position graphs represent the first strike (blue), reverse flow (rf, grey), and backflow (bf, green) phases. The vertical lines in the bone kinematic plots represent the time of the peak gape during the first strike (in blue), peak hyoid depression (in orange), and peak opercula abduction (pink) during the rfs. The sketches on the right represent a lateral and ventral view of a carp (top) and a tilapia (bottom) with the location of the locators used to calculate the gape, hyoid depression, and opercula abduction. Note that the opercula were out of the field view at the beginning of the selected tilapia sequences; therefore, the opercula abduction trace starts after the first strike.

The kinematics during the bf phases corresponded to the same pattern of motions, but with lower magnitudes for the peak gape and hyoid depression than during the intake.

During the rf phase (highlighted in gray in Fig. 5), the sequence of peaks was inverted with respect to the intake: A peak in the opercula opening was followed by a peak in hyoid depression, followed by a peak gape. This reverse sequence of peaks, when compared with the intake, occurred in each trial for both species (Data file S1).

## Discussion

This first application of neutrally buoyant X-ray particle tracking in biological research allowed us to quantify 3D waterflow patterns inside the mouth of a live fish with satisfactory spatial and temporal resolution. Despite the inevitably relatively low number of tracked particles (< 20) compared to common, light-optical techniques using dense suspensions of considerably smaller seeding particles (e.g. particle-tracking velocimetry), importantly for our study, we managed to resolve the flow speeds and directions in the vicinity of the food. Compared to previously employed invasive techniques to quantify intraoral flows, such as endoscopy ^13,25^ or pressure recordings (e.g. ^26^), our non-invasive approach yields a more complete view of the spatial aspects of intraoral suction-feeding dynamics, allowing us to demonstrate the existence of different hydrodynamic mechanisms for food transport.

The main question was how food ends up at the esophagus entrance when the water that carries the food is traveling elsewhere—namely, to the outflow at the gill slits. If water was a pure “carrier” of the food, it would follow our water tracer’s trajectory from the entrance of the buccal cavity to end up in physical contact with the gill rakers (Fig. 1). However, both carps and tilapias appeared to be well capable of positioning the food close to the midsagittal streamlines, thereby avoiding the food from becoming laterally deviated and ending up adhering to the branchial basket. Despite that only a small proportion of the water pathlines passed close to the pharyngeal jaws (Fig. 3), direct interception of the food item by the pharyngeal jaw was common, which avoided the need for subsequent intraoral food-handling actions. Consequently, the efficiency of suction feeding is enhanced by mechanisms that centralize the food paths.

We hypothesize that fish make use of two mechanisms to cause the food to follow a midsagittal trajectory toward the esophagus: (1) developing a ‘central jet’ flow pattern inside the buccal cavity (Fig. 1c), and (2) exploiting the food’s inertia and finite size. Our study found support for the first mechanism by showing a centrally concentrated flow with reverse flow developing near the borders of the buccal cavity (Fig. 4). This central jet (Fig. 1c) was previously reported in computational modelling studies ^21,22^, but was not given any attention thus far. Since a narrowing of the main suction stream will increase the chance of food interception close the esophagus entrance by the pharyngeal jaws (Fig. 1b-c), this will enhance suction efficiently. Further research will be required to unravel why the central jet was considerably more prominent in tilapia (Fig. 4b) compared to carp (Fig. 4a), which should be related to differences in mouth shape, buccal cavity shape, or expansion kinematics.

Secondly, the path curvature from a dorsoventral viewing perspective was significantly higher for the water tracers compared to the food tracers (Fig. 4), indicating that inertial effects are at play. For the large prey that fill the bulk of the buccopharyngeal cavity, such inertial effects would be logical and intuitive. However, our data suggest that these mechanisms, in essence a type of filtration, are also present and effective for a relatively small pellet. A similar mechanism is hypothesized to underlie the process of cross-flow filtration in filter feeders in the flows that pass close to the branchial arches ^12,27^, but here we show that such an inertial separation mechanism is present at the full scale of the buccopharyngeal cavity. The effect of the size, shape, and density of food on its trajectory in a flow field, as shown in our work (Fig. 3), remains to be further quantified to fully understand the constraints of the observed filtration mechanism during the initial phase of food intake. Although flow patterns matching those of a branchial crossflow could not be identified (Fig. 1d), the fish may not have employed this technique as the food type that was offered may not require it to be filtered in such a way.

As shown in previous work ^15,20,28–30^, the feeding process is far from finished after the initial capture in the buccopharyngeal cavity, especially in omnivorous species that perform food sorting and processing. We observed regular and periodical back-and-forth motions of the water tracers after this intake, which moved the food back and forth inside the buccal cavity, with the food finally ending up near the entrance of the esophagus (Fig. 2). Rfs (directed toward the anterior part of the buccal cavity) were previously observed in carps ^13^ and in tilapias ^25^. In carps, anterior movement of the food has been observed using X-ray videos, after which a muscular structure packed with taste buds—the palatal organ—is hypothesized to locally bulge to clamp onto edible particles between the pharyngeal roof and floor, while small waste particles are flushed through the branchial slits ^20,31^. The anteromedial direction of the flows observed in the current study during the rf phase (Fig. 3b) seems ideal to bring food particles from both the gill rakers and from the pharyngeal region toward the palatal organ. Overall, these back-and-forth flows are clearly important in the process of sorting food items from the non-edible particles present in the ingested water, and they potentially also prevent the gill slits from clogging. Rfs most likely involve the gill slit functioning as a flow inlet, a function that has rarely been studied ^32^.

Our data provide the first empirical evidence of bidirectional waterflows being synchronized with motions of the upper and lower jaws, hyoid, and opercula to create anterior-to-posterior waves during bfs and posterior-to-anterior waves during rfs. The anterior-to-posterior wave of head expansion and compression is already well known in fish ^17,32–34^. This is interpreted as a mechanism to both ingest and dynamically accommodate a volume of water as it moves anteroposteriorly though the buccal cavity. For both species studied, this classical anterior-to-posterior wave was also observed during the consecutive periodic bf motions. Yet, the associated waterflow was not purely anterior-to-posteriorly directed: the water was also guided toward the ventral part of the buccal cavity (Fig. 3) by the hyoid depression, and it deviated toward the gill covers (Fig. 4) via the abduction of the opercula (Fig. 5). During the backward motions of the water, we never observed any particles entering the digestive tract, thus consolidating our hypothesis that no water enters inside the esophagus. A wave in the opposite direction was synchronized with the rfs, as hypothesized earlier ^20,32^. The stereotypical bf and rf alternations eventually lead the food to the centerline of the buccal cavity and closer to the pharyngeal jaws, where it will be actively transported toward the digestive tract ^29^.

The new particle-tracking technique we described here (Figs. 6a–d) provided unique data on waterflows during aquatic feeding. It can easily be combined with 3D motion-reconstruction protocols for skeletal elements (i.e. XROMM or X-ray reconstructions of the moving morphology; ^35^) to link waterflow patterns to skeletal kinematics (Fig. 5). The proposed water tracer design had to deal with the conflicting requirements of sufficient radio-opacity while remaining neutrally buoyant, and additionally being small enough to follow small-scale waterflows and avoid interference with the fish’s natural behavior. The easy-to-build protocol (Figs. 6a–d) may be further optimized to reduce the variability in density (Fig. 6c) so that many particles can be kept in suspension in the water column for longer periods. It has strong potential to become an indispensable tool for studying form–function relationships and the biomechanics of various functions involving water motions to which we lack optical access.

**Fig. 6.**
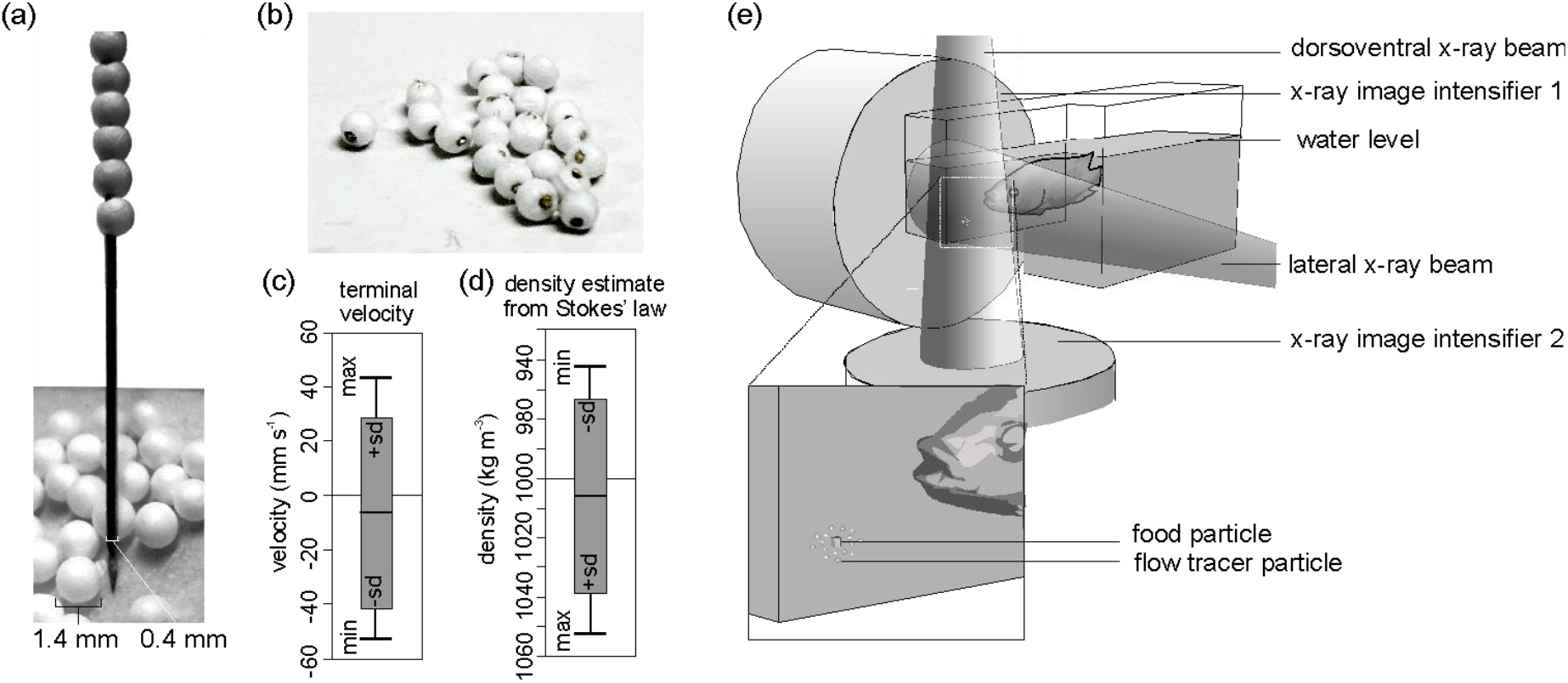
Tracer fabrication for X-ray particle tracking and the experimental design. (**a**) Construction of the flow tracer particles by piercing 1.4-mm-diameter expanded polystyrene (EPS) spheres on a 0.4-mm-diameter, Vaseline-coated brass rod with a sharpened tip. (**b**) Flow tracers after cutting the rod. (**c**) Terminal velocity test results of the EPS brass flow tracers released in water (positive velocities: rising; negative velocities: sinking), and (**d**) the corresponding estimates of density based on Stokes’ law (see materials and methods for details) (*N* = 19). (**e**) Experimental design indicating X-ray beam, image intensifier orientation, and placement of the food surrounded by flow tracer particles at the bottom of the narrow extrusion of the aquarium.

## Materials and Methods

### Animal care and surgical procedures

Two carps (*Cyprinus carpio*), carp01 (mass = 341 g, total length = 25.94 cm) and carp02 (mass = 304.5 g, total length = 20.42 cm) purchased from a pond shop (De Hof-Leverancier, Ekeren, Belgium) and two Nile tilapias (*Oreochromis niloticus*), tilapia01 (mass = 488.4 g, total length = 29.4 cm) and tilapia02 (mass = 489.0 g, total length = 30.3 cm) from a farm (Til-Aqua, Someren, The Netherlands), were housed in two large water tanks at the University of Antwerp, and provided with food ad libitum.

Prior to videography, the fish were anesthetized with 50 mg L^-1^ MS222 (Ethyl 3-aminobenzoate methanesulfonate) and implanted with 0.35-mm-diameter beads made of a tin and silver alloy. Using hypodermic needles, four markers were implanted in the neurocranium, and three on the upper jaw, lower jaw, hyoid, and left operculum (Fig. S1). The left suspensorium, left cleithrum, left fin, and two of the branchial arches were implanted with one or two markers for reference. We also implanted 4-mm-diameter food pellets with three 0.45-mm-diameter markers. After the surgery, the fish were kept under observation until full recovery. The experiments were ethically approved by the University of Antwerp (ECD-2017-22).

### Water tracers

#### Design of the water tracers

The water tracers had to fulfill two typically incompatible requirements: 1) being radioopaque to be trackable on the X-ray videos and 2) being small enough and neutrally buoyant to passively follow the water trajectory as closely as possible. An easy-to-execute and cost-efficient procedure was developed to make spherical waterflow tracers of minimal size to be used with biplanar high-speed X-ray video set-ups. Like the existing tracer designs ^36,37^, we surrounded a radio-opaque metal with a closed-cell foam. The foam compensates for the inevitably high weight of the metal so that the overall density of the tracer particle approximates the density of water.

Expanded polystyrene (EPS) foam spheres with diameters between 1 and 2 mm were purchased (Rovul, Middelstum, The Netherlands) and sieved to retain only those with a diameter of 1.4 mm. To do so, the foam spheres that were loosely stuck in between the mesh holes of a precision test sieve were extracted. Next, 0.4-mm-diameter brass rods (Albion Alloys, Bournemouth, UK) were cut into approximately 0.1-m pieces in length, and their tips were sharpened by holding the rod at a sharp angle against a grinding stone rotating on a Proxxon MF70 micro-milling machine (Proxxon GmbH, Föhren, Germany). Next, the sharp tips were dipped in petroleum jelly to reduce friction. A foam sphere was then gently immobilized between two fingertips while being pierced (as close as possible through its center) by the sharp tip, which was gently rotated by the fingertips of the other hand (Fig. 6a, Movie S3). The one-by-one pierced EPS spheres were slid to the back of the rod. Finally, when the rod was filled with a chain of foam spheres, each sphere was clipped close to the foam sphere edges using side-cutting pliers (Fig. 6b).

Although these particles are large compared to traditional (non-X-ray-based) particles used in particle-tracking velocimetry, the volume of our 1.4-mm-diameter tracers is more than five times smaller than the smallest tracers used so far for X-ray-based particle tracking (^37^; 2 × 2 × 2 mm cubes), which in turn were already considerably smaller than the 8-mm-diameter spheres for which a manufacturing protocol has been described in detail ^36^. In relation to the size of the buccopharyngeal cavity of the specimen studied—roughly 65 × 50 × 40 mm—these tracers are sufficiently small to resolve the large-scale flow structures.

#### Estimation of the food tracer volume and density

To estimate the volume of the food items used in the experiment, we laser-scanned 10 food items (Faro Laser ScanArm V2 system). We obtained a volume (mean ± sd; N = 10) of 24 ± mm^3^.

The food items implanted with radio-opaque markers were weighted, to obtain their dry mass and were soaked in water for 15 minutes to estimate their wet mass. We obtained a density (mean ± sd; N = 10) of 1845 ± 170 kg/m^3^.

#### Test of the water tracer density

Nineteen water tracers held by forceps in the middle of a water column were released and filmed at 50 frames per second next to a scale bar to extract their trajectory using XMALab software version 1.5.1 (www.xromm.org/xmalab/) ^38^. We measured the terminal velocity (V_terminal_) of the tracer (Fig. 2c) and calculated the water tracer density (ρ_tracer_) using Stokes’ law (Eq 1):

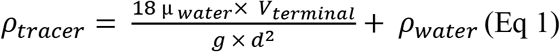

where μ_water_ corresponds to the dynamic viscosity of water—that is, 1.002 mPa.s at 20°C^39^—, g to the gravitational acceleration of 9.81 m.s^-2^, d to the diameter of the tracer including both the nickel rod and the foam sphere—that is, 1.4 mm—, and ρ_water_ to the water density of 1000 kg.m^-3^. The mean (± standard deviation) density of the produced particles from the test sample (*N* = 19) was 994 ± 33 kg.m^-3^ (Fig. 6d). Since only negatively buoyant particles were presented at the bottom of the aquarium surrounding the food (Fig. 6e), the mean particle density was 1031 ± 14 kg.m^-3^, with a maximum of 1049 kg.m^-3^ (*N* = 11).

#### Biplanar X-ray videography

The four implanted fish were recorded during their feeding on a marked food item surrounded by about 15 radio-opaque water tracers. They were filmed at a resolution of 2048 × 2048 pixels and at a speed of 750 frames per second using the 3-D DYnamic MOrphology using X-rays (3D^2^YMOX) set-up at the University of Antwerp ^40^. The system consists of two X-ray videography systems, each composed of a Photron FastCam Mini WX50-32GB camera (Photron USA, Inc., San Diego, CA, USA) mounted on a Philips Imagica HC 38-cm image intensifier. The two conical X-ray beams were set to create a lateral and a dorsoventral view (Fig. 6e) using the image intensifier’s smallest field-of-view setting for the carp and the medium field-of-view setting for the tilapia, emitting at tube voltages and currents of 75 kV and 80 mA and 95 kV and 85 mA, respectively (Fig. 6e). More technical details and the performance tests of the 3D^2^YMOX system can be found in a previous publication ^40^.

We followed recommendations regarding distortion correction and space calibration during the experiment ^35,41^ and used a perforated steel sheet with a precise hole size and spacing for distortion correction and a calibration object for determining camera 3D-view characteristics. We recorded 73 sequences of suction feeding of around 5 seconds in duration and selected 15 sequences corresponding to successful attempts (e.g., fish head in the field of view during the entire sequence, water tracers successfully ingested with the food item) balanced among individuals. The fish were euthanized with MS222 and scanned using microcomputed tomography (μCT), with the implanted markers intact (μCT scans are available at the Royal Belgian Institute of Natural Sciences; 110 kV, 0.5 mA and a resolution of 64.94 μm on each axis). We used Avizo (version 6.3; FEI Visualization Sciences Group) to reconstruct bone models and the implanted markers from the μCT scans.

### X-ray reconstruction of the moving morphology

Using the validated XMALab software version 1.5.1 (www.xromm.org/xmalab/)^38^, we undistorted the video images, calibrated the view of the X-ray cameras, and tracked the three types of markers (bone markers, food and water tracers) on each frame of the selected videos.

#### Rigid bodies

To reconstruct the 3D motion of the implanted bones, we followed the XROMM workflow; we imported the 3D bone models into Maya (version 2018; Autodesk) and extracted CT marker coordinates. We generated the rigid body transformations in XMALab and imported them into Autodesk Maya to animate the bones. We used scientific rotoscoping ^41^ to position rigid bodies with less than three markers (in case of losses after surgery or when markers were out of view).

#### Coordinate system definition

To obtain a common framework between trials, we created an anatomical coordinate system (ACS) attached to the neurocranium. The origin of this ACS was offset to be positioned close to the location of the pharyngeal jaws as an approximation of the esophagus entrance location (midsagittal plane, caudal tip of the chewing pad). The pharyngeal jaws are known to move during the food capture and transport (e.g., ^20,42^), but we did not reconstruct their motions in the present study. Their static position was used as an estimation of the esophagus entrance location. The x axis was parallel to the plate of the pharyngeal process, pointing toward the entrance of the buccal cavity, and was positive rostrally. The y axis corresponded to the midsagittal axis, perpendicular to the x axis, and was positive dorsally. The z-axis vector (positive right) was calculated by crossing the proximal x axis with the y axis (Fig. 2).

#### Food and water tracers

We imported the 3D trajectories of the food and water tracers into Maya (version 2018; Autodesk). We computed the relative 3D trajectory and relative 3D velocity in the coordinate system previously described for the water and food tracers (Fig. 2, Movie S3).

#### Phases of suction feeding

To divide the suction-feeding sequences into phases, we used the anteroposterior trajectory and velocity of the water tracers (Data file S1). We excluded the stagnant and slow-moving parts of the path lines by discarding velocities between −4.5 and 2 cm/s from the analysis.

The 3D trajectory of all the water tracers during each suction-feeding sequence provided a visualization of the water tracer paths for the different views of a representative trial and of all the trials of a given species during the intake (Fig. 3a), the rf phase (Fig. 3b), and the bf phase (Fig. 3c).

We registered the number of successful suction-feeding sequences among all the recorded trials (*N* = 37 in carps, *N* = 36 in tilapias). For this set of data, we noted the number of food and water tracers reaching the entrance of the esophagus that were effectively trapped in the digestive tube when the tracer was at the level of the pharyngeal jaws. We also measured the number of water tracer oscillations (backward and forward motions) before the food tracer reached the area of the pharyngeal jaw’s axis.

In addition, we compared the global distance traveled by the food tracers and the water tracers (Fig. S2). We computed the relative contribution of each component of the trajectory during the intake, the rf phases, and bf the phases among the studied trials (Table S1).

### Path line curvature

To quantify the deviation of the water tracers relative to the food tracers, we calculated the curvature of a given tracer based on three consecutive frames in the dorsoventral plane (x, z). To do so, we standardized the position of the tracers on the anteroposterior axis by calculating the x-trajectory percentage for each tracer, then we computed the curvature values over these intervals for the food tracers and the water tracers. As the distribution was not normal, we performed a Kruskal–Wallis rank sum test to test if the differences in the maximum curvature per trajectory of each interval were significantly different between the water tracers and the food tracers (Fig. 4, Data file S1).

### Skeletal kinematics

We exported the 3D trajectories of additional locators we positioned on the anterior tip of the upper jaw, lower jaw, and hyoid, as well as on the posterior tip of the operculum on the animated Maya scenes (Fig. S1). This allowed us to compute the gape, hyoid depression, and opercula abduction for six carp sequences and three tilapia sequences. We extracted the time when the peaks of each variable were reached during the intake and the rfs for each analyzed sequence.

## General

We would like to thank Peter Aerts for access to the facility at the University of Antwerp, Jonathan Brecko for the CT-scan acquisition at the Royal Belgian Institute of Natural Science, as well as Anthony Herrel for helpful discussions.

## Author contributions

PP and SVW designed the experiment and acquired the experimental data.

PP, AB, and AF analyzed the data. PP and SVW wrote a first draft of the manuscript, which was read and edited by all the authors.

## Supplementary Materials

**Fig. S1.**
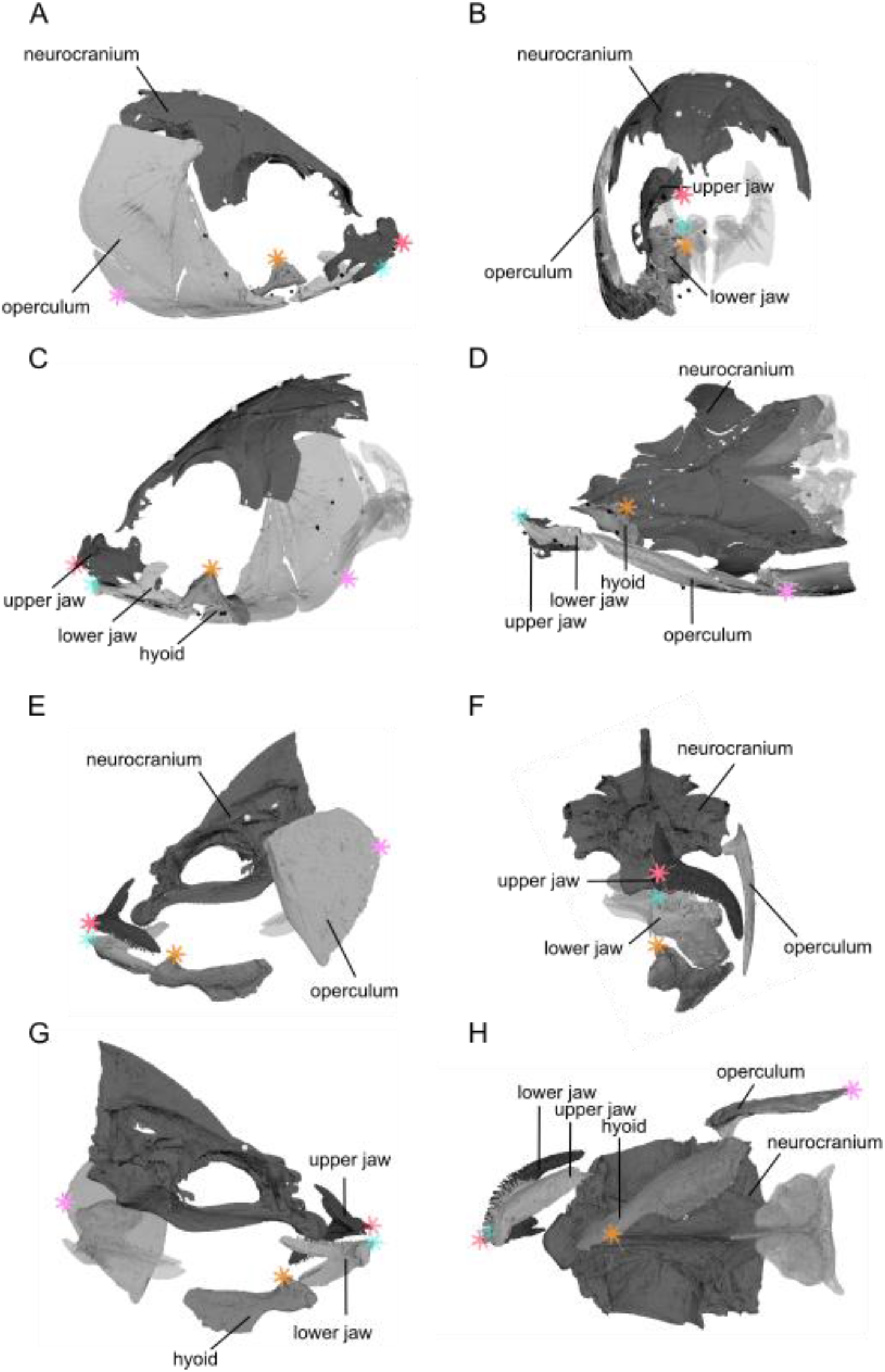
Position of the implanted markers and the locators in the carps (A, B, C, and D) and tilapias (E, F, G, H). The lateral views (**A, C, E, G**), frontal views (**B, F**), and dorsal views (**D, H**). The pharyngeal jaws are transparent, the implanted markers are in black, and the locators added to the rigid bodies are in color.

**Fig. S2.**
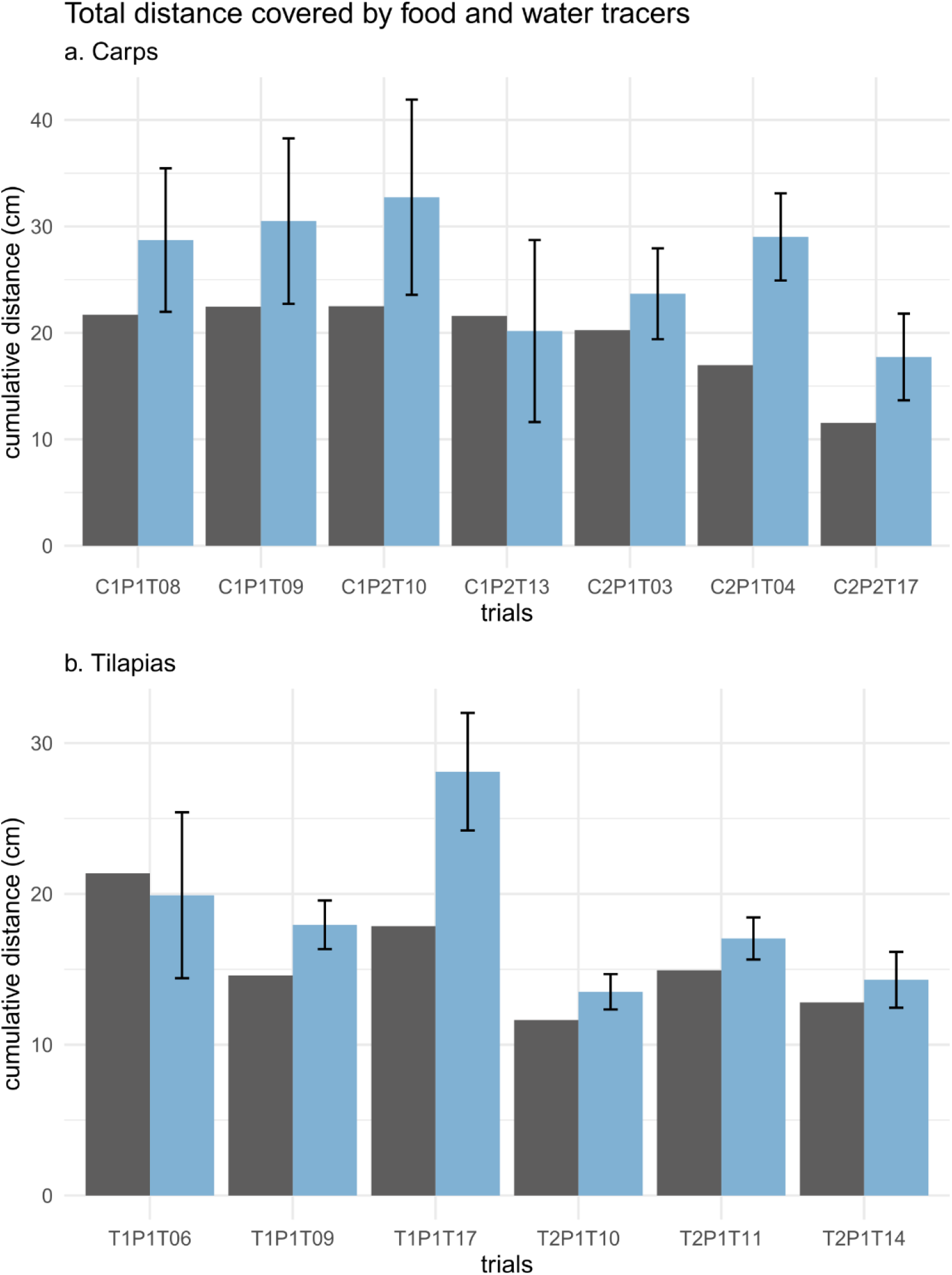
Histograms of the total distance covered by the food (in black) and the water tracers (in blue) during an entire sequence of suction feeding for each carp (a) and tilapia (b) trial.

**Table S1.** Normalized displacement components along the three cranium-bound orthogonal axes shown in Fig. 1 (mean ± SD in %) for the food and water tracers in carps and tilapias.

|  | Carps |  | Tilapias |  |
| --- | --- | --- | --- | --- |
|  | food | water | food | water |
| **Intake** |  |  |  |  |
| Anteroposterior (x) | 64 ± 1 | 64 ± 1 | 57 ± 1 | 63 ± 0.1 |
| Dorsoventral (y) | 26 ± 1 | 28 ± 1 | 20 ± 0.4 | 14 ± 0.3 |
| Lateromedial (z) | 10 ± 1 | 8 ± 0.4 | 23 ± 0.1 | 22 ± 0.1 |
| **Reverse flow** |  |  |  |  |
| Anteroposterior (x) | 52 ± 1 | 54 ± 1 | 39 ± 1 | 46 ± 3 |
| Dorsoventral (y) | 29 ± 1 | 30 ± 1 | 19 ± 1 | 20 ± 1 |
| Lateromedial (z) | 19 ± 1 | 16 ± 1 | 42 ± 1 | 34 ± 2 |
| **Backward flow** |  |  |  |  |
| Anteroposterior (x) | 53 ± 1 | 58 ± 1 | 43 ± 1 | 51 ± 2 |
| Dorsoventral (y) | 29 ± 0.5 | 27 ± 1 | 23 ± 1 | 22 ± 1 |
| Lateromedial (z) | 18 ± 1 | 15 ± 1 | 34 ± 1 | 27 ± 2 |

---

**Movie S1: Video of a suction-feeding sequence of a carp, showing the raw video, the water and food tracer trajectories, and the rigid body reconstruction.**

**Movie S2: Video of a suction-feeding sequence of a tilapia, showing the raw video, the water and food tracer trajectories, and the rigid body reconstruction.**

**Movie S3: Video summarizing the present study.**

**Data file S1: Vignettes coming from the R package associated with the building of the data IOFLOW package, giving an overview of the methods used to generate the data (1. Clean Raw Data) and the analysis presented in the manuscript (2. Analyze Cleaned Data).**

